# Optimization of *Agrobacterium*-mediated transformation of commercial heirloom tomato cultivars to develop novel traits via CRISPR/Cas9 Genome Editing

**DOI:** 10.64898/2026.02.07.703897

**Authors:** Jordan Oxendine, Elsa Ibarra-Reyes, Junchi Ma, Carrie Li, Sarah Baron, Allison E. Hwang, Ruiting Wang, Daniel Rodriguez-Leal

**Affiliations:** Tomato Lab. Department of Plant Science and Landscape Architecture. University of Maryland, College Park, MD, 20742, US

**Author notes:** **Author information:** Equal contribution (Jordan Oxendine, Elsa Ibarra-Reyes). **Main conclusion** Trait development for commercial heirloom tomatoes can be advanced by optimization of tissue culture and transformation via Agrobacterium and CRISPR/Cas9 mutagenesis.

**Keywords:** Trait development, Biotechnology, Genetics, Tissue culture

## Abstract

Genetic improvement using new genome editing approaches rely on the efficient delivery of the CRISPR/Cas system in the vegetable crop tomato. Previous protocols for tomato transformation have primarily focused on a handful of cultivars (M82, Alisa Craig, Microtom, Sweet-100) with very little commercial relevance, and it is not clear if these protocols can be implemented directly in other commercially relevant varieties. Heirloom tomatoes are sought for their deep and diverse flavor but have not been subjected to systematic crop improvement via conventional breeding or biotechnology approaches such as transgenesis or genome editing. Therefore, we tested the transformation and regeneration capacity of six different heirloom cultivars known for their superior taste and market relevance in the US. Subsequently, we optimized rooting conditions and used the *GRF4-GIF1* chimeric developmental regulator to successfully recover transgenic plants. Finally, we evaluated the efficiency of targeted genetic modification using the CRISPR/Cas9 genome editing system in several of these cultivars. We demonstrate that our optimizations led to successful transformation of several heirloom varieties, including the generation of edited plants for target genes modifying plant architecture and flowering time. Our results set the foundation for a biotechnology platform to deliver improved traits to local and regional heirloom varieties using genome editing.

## Introduction

The vegetable tomato (*Solanum lycopersicum*, L.) is among the most widely cultivated and consumed vegetables globally (Govindasamy et al 2025;FAOSTAT 2023). Global tomato production reached approximately 190 million tons in recent years (OECD 2017; FAOSTAT 2023). Besides their global relevance, tomatoes are also important for local and regional consumption, particularly in medium-scale, controlled-environment and urban agriculture settings (USDA ERS 2024). Heirloom tomatoes are known for their diverse and flavorful fruit types (Ronga et al. 2021; Dwivedi et al. 2019; Marcinkiewicz 2017). These cultivars are old (≥50 years) varieties that have been passed onto generations as seeds from open pollinated plants grown in gardens or fields from local farmers. Heirloom tomatoes are sought for their deep and diverse flavor, but have not been subjected to systematic crop improvement via conventional breeding or biotechnology approaches such as transgenesis or genome editing. Improvement of these varieties could lead to increased yield potential and better return-on-investment for small-to-medium scale farming businesses and for seed companies investing in local and urban markets.

Tomato is also considered a crop model used for genetic and developmental studies to understand gene function in plant development and crop improvement (Wang et al. 2024), including the implementation of genetic engineering via plant transformation and regeneration. Several studies have shown that tomato is relatively amenable to tissue culture and plant transformation using *Agrobacterium tumefaciens* (Agrobacterium) (Frary & Earle 1996; (Gerszberg et al. 2014; Kaul et al. 2024). The efficiency of Agrobacterium transformation is determined by the type and age of explants, the bacterial strain used, and the nutrients and growth regulators used in the media during co-cultivation and regeneration (Frary & Earle 1996; Gerszberg et al. 2014). Notably, previous protocols for tomato transformation have primarily focused on a handful of cultivars (M82, Alisa Craig, Microtom, Sweet-100) with very little commercial relevance. Although these protocols could be used, it is likely that optimizations would be required for successful transformation of commercially-relevant cultivars such as heirlooms.

In order to apply novel crop improvement technologies such as genome editing in cultivars with commercial relevance for local, national and international markets, it is critical to develop efficient and consistent protocols for plant transformation and regeneration. To address this gap, we have implemented Agrobacterium-mediated transformation in several heirloom varieties known for their superior taste and local/regional relevance in the US market. We first tested a known protocol for regeneration (Van Eck et al. 2019) to evaluate its efficiency in six different heirloom cultivars. Subsequently, we optimized the protocol by adjusting the media and conditions during the rooting stage, and by using the *GRF4-GIF1* chimeric gene to enhance regeneration and successfully recover transgenic heirlooms. Finally, we evaluated the efficiency of targeted genetic modification using the CRISPR/Cas9 genome editing system in several of these cultivars. We demonstrate that our optimizations led to successful transformation of six heirloom varieties, including the generation of edited plants for target genes modifying plant architecture and flowering time. Our results set the foundation for a biotechnology platform to deliver improved traits to local and regional heirloom varieties using genome editing.

## Materials and Methods

### Plant Material and growth conditions

Tomato seeds were surface sterilized with 70% ethanol for 30 seconds and sodium hypochlorite (2%) solution for 10 min, washed thoroughly with sterile distilled water five to six times and placed on half-strength Murashige and Skoog (Murashige and Skoog, 1962) medium for germination in small glass jars. Seeds were grown under long-day conditions (16-h light/ 8-h dark) in a growth chamber (Percival Scientific) supplemented with artificial white light from LED panels (~200 umol/m^2^) at 25°C and 50% humidity. Seeds from M82 were kindly donated by Prof. Zachary Lippman at CSHL. Seeds from Heirloom varieties were purchased from commercial vendors (Johnny’s Seeds and Annie’s heirloom seeds). For greenhouse experiments, seeds were directly sown in soil in 96-cell plastic flats. Plants were grown in a greenhouse under long-day conditions (16-h light/8-h dark) supplemented with artificial light from high-pressure sodium bulbs (250 µmol/m^2^).

### Guide RNA (gRNA) designs for CRISPR/Cas9 gene editing of *Brachytic* and *SP5G*

For CRISPR/Cas9 constructs, we selected previously-published gRNA designs (Soyk et al. 2017; Lee et al. 2022) targeting the coding sequences of *Br* (*Solyc01g066980*) and *SP5G* (*Solyc05g053850*), respectively. The specificity of these gRNA was confirmed by BLAST against the most recent version of the Heinz genome (SL4.0) in Solgenomics. Subsequently, we synthesized these gRNAs as primers through an external vendor (Azenta).

### Vector construction

Vectors for plant transformation were assembled using the MoClo Golden Gate cloning system as previously described (**Fig. S1**) (Werner et al. 2012). First, the *NPTII* gene used for selection during tissue culture was sub-cloned from pICSL7004 (addgene plasmid #50334) into a Level 1 (L1) vector (pICH47732) to make pTL0038. This L1 vector was subsequently cloned along with pICH47742-35S:Cas9 (Addgene plasmid #49771) into the binary backbone pAGM4723 to make the Level 2 (L2) vector pTL0047. To produce each gRNA L1 vector, a PCR reaction was carried out with a primer containing the gRNA sequence (**Table S1**), using the plasmid pICH86966:: AtU6p::gRNA_PDS (Addgene plasmid #46966) as the template. Each gRNA was cloned individually into the L1 vectors in forward orientation. To assemble the L2 vector pTL0151 we combined pTL0038, pICH47742-35S:Cas9 and the gRNAs with the binary L2 vector pAGM4723. To create pTL0153, we combined the pTL0038 with an L1 vector containing the *GRF4-GIF1* chimeric gene under the regulation of the parsley *UBIQUITIN* promoter and the Pea *3A* terminator (gift from Professor Sebastian Soyk, University of Lausanne) with pICH47742-35S:Cas9, and the *Br* and *SP5G* L1 gRNA vectors. All vectors were confirmed by enzyme restriction and whole-plasmid sequencing through an external vendor (Azenta).

### *Agrobacterium*-mediated plant transformation

*Agrobacterium tumefaciens* (Agrobacterium) strain GV3101 was used for all experiments. All binary vectors were introduced into the competent cells through the freeze-thaw method (Weigel and Glazebrook 2006). A single colony was taken from LB-plates containing kanamycin, rifampicin and gentamicin at 50 mg/L each, and grown in 15 ml of LB medium containing 50mg/l Kan at 28°C with constant shaking (220 rpm) for 16 hours. The bacterial suspension was centrifuged and resuspended in MS-0.2% medium (4.3 g/l MS salts, 2% sucrose, 100 mg/l myo-inositol, pH 5.8) to optical density (0D_600_) of 0.6, and used for explant infections.

Agrobacterium-mediated infections were performed following a previous protocol (Van Eck et al. 2019) with minor modifications. The cotyledons from seedlings 8 days after germination (8 DAG) were dissected into two squared sections each (explants) and incubated on 2Z-medium (4.3 g/l MS salts, 2% sucrose, 100 mg/l myo-inositol) at 25°C for 24 h before transformation. Explants were co-cultivated with Agrobacterium on 2Z-medium for 48 h at 25°C in the dark and then transferred to 2Z selection medium supplemented with 200 mg/L Kan. Explants were cultivated in a growth chamber under white LEDs (16h light/8 h dark, 200 µmol/m^2^) at 25°C and 50% humidity and transferred every two weeks to fresh selection media until visible shoots appeared. Shoots were excised and transferred into plastic containers with selective rooting medium supplemented with 200 mg/L Kan, 0.5mg/L Indole-3-butyric acid (IBA) and 0.5 g/L activated charcoal (Ludwig-Müller et al. 2005; Thomas 2008). Well-rooted shoots were transplanted to soil and acclimated in a greenhouse environment with day temperature of 24-28°C and 60-80% relative humidity.

### Molecular Characterization of edits generated by the CRISPR/Cas9 system

Tomato genomic DNA was extracted from putative transgenic tomato leaves by using a variation of a fast-track DNA extraction protocol (Guo et al. 2022) that included the addition of a chloroform:isoamyl alcohol (24:1) extraction instead of using 5M Potassium Acetate. Briefly, we collected the tips of the cotyledons from growing seedlings into 96-well plates along with a 4 mm steel bead. Tissue samples were ground in a Genogrinder (HG-600, Cole Palmer), followed by the addition of 350µL of lysis buffer and 50µL of 10% SDS. Samples were then incubated in an oven at 65°C for 25 min. Subsequently, 400µL of chloroform:isoamyl alcohol (24:1) were added to each sample, vortexed vigorously and centrifuged at 4,000 rpm for 25 min. We recovered 120 µL of the upper phase from each sample into a 96-well PCR plate and mixed it with an equal volume of isopropanol. Samples were then centrifuged at 4,000 rpm for 25 min, followed by a washing step with 70% Ethanol and subsequent centrifugation at 4,000 rpm for 10 min. Samples were then air-dried for 10-20 mins and eluted in DNAse-free water supplemented with 0.05 µL of RNAse (10mg/mL, Thermo Scientific) per each 50µL of water. Plates were incubated at 65°C for 30 mins, followed by a quick vortexing and centrifugation (1 min at 4,000 rpm) and kept at −20°C until use. To perform genotyping for the presence of the transgene and the edits, we used specific primers matching the target regions of *Br* and *SP5G* **(Table S2)**. PCR was carried out to amplify specific fragments from putative transformed plants and their progeny using a RedTaq Polymerase (Genesee Scientific). The approximate product lengths of the transgene target site, *SP5G* and *Br* were 500, 400, 300 bp respectively. The PCR program used was: 1 cycle of initial denaturation at 95°C for 3 minutes, followed by 30 cycles of 95°C for 30s, 72°C for 30s and 72°C for 30s. The amplified products were analysed by gel electrophoresis in 1% (w/v) agarose gels and visualized in a gel imager (Nippon Genetics).

## Results

### Initial evaluation of transformation efficiency in multiple tomato cultivars

To determine the transformation efficiency and regeneration potential of multiple heirloom cultivars commercialized in the Mid-Atlantic US, we initially implemented a previous protocol successfully used in the cultivar M82 (Van Eck et al. 2019). This protocol uses cotyledons from germinated seedlings as explants for co-cultivation with Agrobacterium. Under our growth conditions, most cultivars germinated within 7-11 days. We used seedlings with expanded cotyledons without visible true leaves. Using a limited number of explants per cultivar, we performed an initial transformation using the Agrobacterium strain *GV3101* carrying the plasmid pTL0047 containing the selectable marker gene *neomycin phosphotransferase II* (*nptII*), allowing for selection of putative transgenic plants using kanamycin. Our initial evaluation consisted of the heirloom cultivars Jubilee, Amana Orange, Sunray, Brandywine Pink, Red potato leaf Brandywine, Mortgage Lifter and the reference processing tomato vintage cultivar M82. These heirloom varieties represent different fruit types (round, small or beefsteak, red, brown or yellow color fruits) and exhibit different maturities (**Table S3**). With this protocol, we observed callus formation at week 4 and emerging shoots at week 6 (**Fig. 1a-b)**. Elongated shoots were transferred into a glass jar with root-inducing media at week 8-10 and then transferred to soil at weeks 10-12. In this first experiment, M82 exhibited ~30% transformation efficiency (number of shoots growing in Kan per total number of explants) (**Fig. 1c**). Among the tested heirloom cultivars, Brandywine Pink and Jubilee were the most recalcitrant (8.8% and 3.4% shoot regeneration, respectively). Notably, while M82 shoots started developing roots within 2 weeks of transferring to root media, most heirlooms showed no signs of rooting after 6 weeks (**Table S4**) in rooting media. These results suggest that heirloom cultivars exhibit differences in regeneration potential compared to M82. In order to recover transgenics from these cultivars, we decided to 1) optimize rooting media and 2) use the chimeric *GRF4-GIF1* developmental regulator for enhancing regeneration.

**Figure 1.**
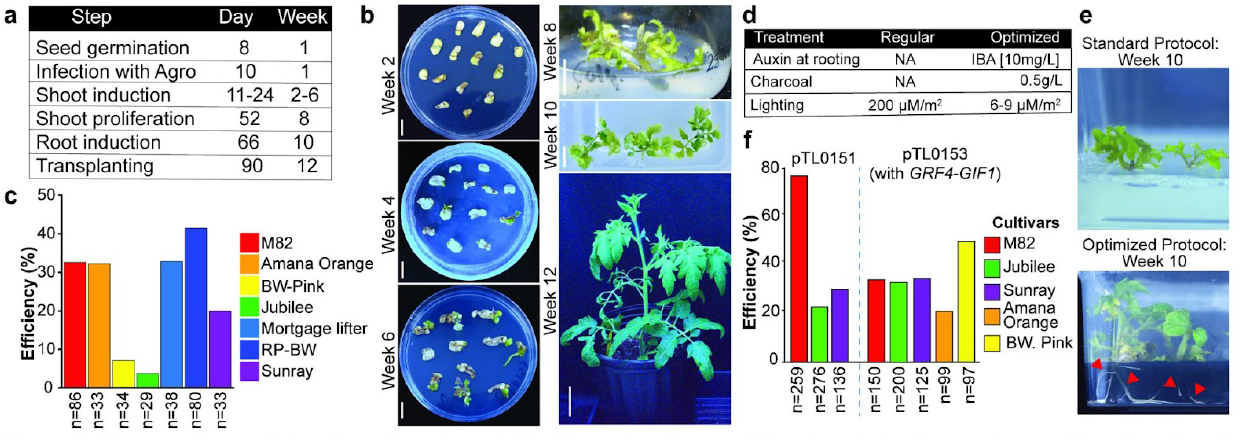
Tissue culture and transformation of several heirloom tomatoes. **a**, Main steps during tissue culture and transformation. **b**, representative picture of each of the stages from **a. c**, Transformation efficiency evaluated based on the number of proliferating shoots over the total number of explants used. **d-e**, To induce successful rooting, different variables were adjusted during the rooting stage, leading to root induction in all heirloom cultivars tested. **f**, Transformation efficiencies in different heirloom cultivars transformed with two different binary vectors (pTL0151 and pTL0153). Scale bars in **b**, 1 cm in plates, 1 cm in week 8 and 10 and 5 cm at week 12

### Optimization of tissue culture conditions to develop genome edited heirloom tomatoes

Despite we observed shoot induction in media containing Kanamycin (200mg/L), all shoots from heirlooms transferred to rooting media failed to develop roots. In our initial experiment, all shoots were transferred to root media without any plant-growth regulators and under the same light conditions (200µm/s^2^) as during the shoot-induction stage. Therefore, we adjusted the regeneration protocol by modifying rooting media and light intensity (**Fig. 1d**). We adjusted the rooting media by adding 0.5mg/L of Indole-3-butyric acid (IBA) and 0.5g/L of activated charcoal. For this second experiment we transformed M82, Sunray and Jubilee with the binary vector pTL0151 (**Fig. 1f**) containing the selectable marker *nptII* and the CRISPR/Cas9 system encoding four gRNAs targeting the plant height and flowering time regulators genes *Br* and *SP5G*. Similar to our first experiment, we observed shooting in week 6 for these 3 cultivars. At week 8, we transferred the excised shoots into plastic containers with rooting media containing IBA and activated charcoal. The plastic containers were placed in a plant-growth rack with ambient lighting (6-9 µmol/s^2^) at room temperature (22-25°C). Notably, we observed the majority of shoots started developing roots in all cultivars within 7 days (**Fig. 1e**). Our results showed the three cultivars exhibited higher transformation and regeneration, with M82 showing the highest efficiency (76.6%) and with Jubilee showing an increase compared to our initial experiment (3.4% vs 22.7%). To characterize editing efficiency, we performed PCR to amplify the target regions matching the *Br* and *SP5G* gRNAs in T0 plants from all three cultivars (**Fig. 2a-b**). Several T0 plants exhibited more than one band suggesting the CRISPR/Cas9 system was creating dropout deletions in the target region of both genes (**Fig. 2c and Fig. S2**). To determine the specific mutant alleles generated in these T0 plants, we performed direct Sanger sequencing on purified PCR products from selected T0 plants from M82, Amana and Jubilee, and analyzed the data using ICE Analysis from EditCo. We found T0s for all three cultivars showing indels and large dropouts in both target genes (**Fig. 2c**). Notably, one of the Amana

**Figure 2.**
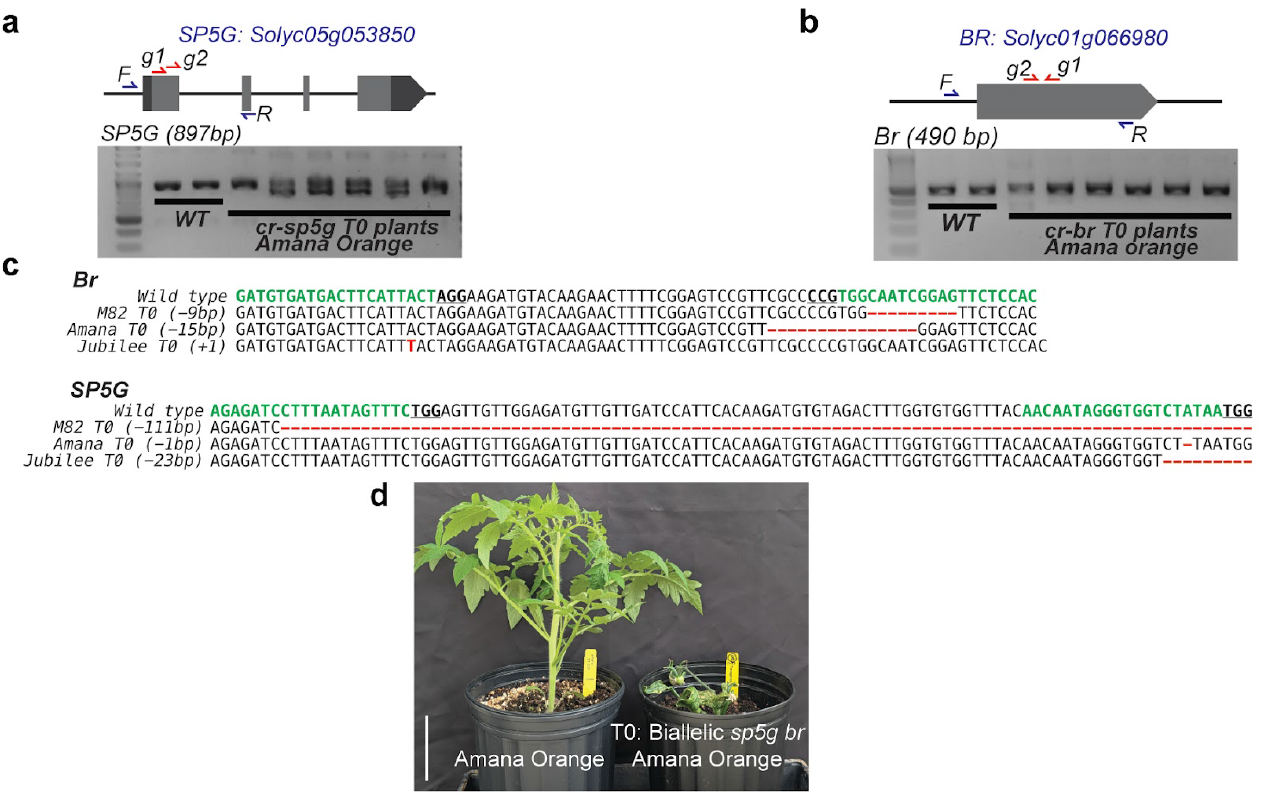
Genome editing using CRISPR/Cas9 in different heirloom tomato cultivars. **a-b**, Gene models and gRNA designs for *Br* and *SP5G* target genes. Green text, gRNA; bold and underlined text, PAM; red text, indels. **c**, Sequencing data on selected T0 showing editing of both target genes in M82, Amana Orange and Jubilee. **d**, Representative picture of a biallelic mutant recovered at T0 stage in the heirloom variety Amana Orange. Scale bar in **d**, 5 cm

Orange biallelic T0 plants was dwarf and early flowering, suggesting the mutations are likely causing complete knockout of both *Br* and *SP5G* (**Fig. 2d**). Our results demonstrate that heirloom varieties required additional optimization of regeneration conditions to successfully produce transgenic and edited plants carrying mutations generated by the CRISPR/Cas9 system.

### Implementation of developmental regulators for regeneration in heirloom tomatoes

Previous studies in different crops suggest the use of developmental regulators (DR) can dramatically increase the rate of transformation in recalcitrant species by enhancing regeneration under tissue culture conditions (Lian et al. 2022). In tomato, a previous report suggested the use of the *GRF4-GIF1* chimera enhanced regeneration in the cultivar Sweet-100 and in a swiss heirloom variety (Swinnen et al. 2025). To test if the *GRF4-GIF1* chimera could enhance regeneration in multiple heirloom tomatoes, we performed a third experiment with a new vector (pTL0153) carrying *nptII*, the CRISPR/Cas9 system targeting *Br* and *SP5G* and a *GRF4-GIF1* expression cassette (**see Materials and Methods**). This time, we expanded our list to include heirloom cultivars used in our first experiment and maintained our improvements during the rooting stage at week 10. We observed that most of the heirlooms varieties transformed with pTL0153 exhibited similar regeneration rooting compared to the initial experiment (**Table S4**). Although we recovered transgenic plants 1-3 weeks earlier for most cultivars transformed with pTL0153, transformation efficiency was only significantly different for cultivars Jubilee compared to transformations using pTL0151 (Two-Proportion Z-Test p-value < 0.01, **Table S5**). These results indicate that *GRF4-GIF1* speeds up the process of shooting and regeneration but does not increase transformation efficiency in all cultivars tested (**Table S6**).

Collectively, our results suggest that heirloom tomatoes are suitable for plant transformation and CRISPR/Cas9-mediated genome editing and that further optimization and the implementation of DRs allow for faster recovery of stable transgenic and edited heirlooms. We found the addition of IBA and activated charcoal ensured the development of roots in the transgenic shoots and allowed for recovery of edited plants showing modified plant architecture. Our work sets the foundation for a biotechnology platform to deliver improved traits to local and regional heirloom varieties using genome editing.

## Data availability

Additional data is provided in the supplemental materials. Seeds of edited heirlooms and vectors designed in this study are available upon a valid request to the corresponding author and following proper material transfer agreements.

## Acknowledgements

We thank all members of the Tomato Lab for their support on this research project. We thank Sydney Wallace, Meghan Fisher, and Christian Mckay for their assistance in greenhouse plant care. We thank Drs. Zach Lippman and Sebastian Soyk for sharing materials. This research was supported by grants from MAES (MD-PSLA-243084) and the California Tomato Research Institute (CTRI), and startup funds from UMD-PSLA.

## Author contributions

J.O., E.I-R. performed tissue culture and plant transformation experiments. J.M., C.L., S.B.,A.E.H and R.W. performed vector construction and genotyping of edited plants. D.R-L. conceived the project, supervised all experiments, performed greenhouse care, genotyping and data analysis, wrote and edited the manuscript with support from J.O and E.I-R. All authors read and approved the manuscript.

