## Supplemental Figures and Tables for "Optimization of *Agrobacterium*-mediated transformation of commercial heirloom tomato cultivars to develop novel traits via CRISPR/Cas9 Genome Editing"

**Title:**

**Author information:**

[1]Equal contribution

[2]Tomato Lab. Department of Plant Science and Landscape Architecture. University of Maryland, College Park, MD, 20742, US.

**Correspondence:**

**ORCiD:** https://orcid.org/0000-0003-1871-7432

| Table S1. guide RNA sequences used in this study | |
|---|---|
| **gRNA id** | **gRNA sequence** |
| *SPG5-g1* | AGAGATCCTTTAATAGTTTC |
| *SPG5-g2* | AACAATAGGGTGGTCTATAA |
| *Br_g1* | GTGGAGAACTCCGATTGCCA |
| *Br_g2* | GATGTGATGACTTCATTACT |

| Table S2. Primers used in this study | | |
|---|---|---|
| **primer id** | **sequence (5´to 3´)** | **purpose** |
| TL0231 | CATTGCGATTCTTGACGACGA | Br genotyping and sequencing |
| TL0143 | GGGAGACTACCCCACTATCCA | Br genotyping and sequencing |
| TL0144 | TTCACGACTTGTCAACCATTG | SP5G genotyping and sequencing |
| TL0199 | GGTTGCTAGGGTTTGGAGCA | SP5G genotyping and sequencing |
| TL0246 | tgtggtctcaATTGAGAGATCCTTTAATAGTTTCgttttagagctagaaatagcaag | SP5G-1 gRNA cloning |
| TL0247 | tgtggtctcaATTGAACAATAGGGTGGTCTATAAgttttagagctagaaatagcaag | SP5G-2 gRNA cloning |
| TL0248 | tgtggtctcaATTGGTGGAGAACTCCGATTGCCAgttttagagctagaaatagcaag | Br-1 gRNA cloning |
| TL0249 | tgtggtctcaATTGGATGTGATGACTTCATTACTgttttagagctagaaatagcaag | Br-2 gRNA cloning |
| TL0250 | tgtggtctcaAGCGTAATGCCAACTTTGTAC | gRNA cloning (universal primer) |

**Table S3. Information on heirloom cultivars used in this study**

| Variety | Days to Maturity | Fruit Color | Fruit Size / Shape | Growth Habit | Notable Characteristics |
|---|---|---|---|---|---|
| M82 | 75–80 | Red | Medium, round | Determinate | Standard research cultivar with strong uniformity. |
| Amana Orange | 85–90 | Deep orange | Very large, beefsteak | Indeterminate | Mild, sweet, low-acid; large fruit set. |
| Brandywine Pink | 85–100 | Pink-red | Very large, beefsteak | Indeterminate | Classic Brandywine flavor; potato-leaf trait. |
| Jubilee | 70–80 | Red-orange | Medium, elongated plum | Indeterminate | Sweet, low-acid; consistent coloration. |
| Mortgage Lifter | 80–85 | Pinkish-red | Large, beefsteak | Indeterminate | High-yielding; rich balanced flavor. |
| Red Potato Leaf Brandywine | 80–95 | Red | Large, beefsteak | Indeterminate | Red-fruited potato-leaf Brandywine type. |
| Sunray | 75–80 | Orange | Medium, round | Indeterminate | Smooth, uniform fruits. |

**Table S4. Raw data on tissue culture and transformation experiments**

| Vector | Cultivar | Total # seeds | Total # explants | Total Infected[a] | Total # of shoots | Regeneration %[b] | Total # rooted | Total in soil |
|---|---|---|---|---|---|---|---|---|
| pTL0047 | M82 | 150 | 86 | 86 | 29 | 33.70% | 35 | 0 |
| pTL0047 | Amana Orange | 150 | 33 | 33 | 11 | 33.30% | 1 | 0 |
| pTL0047 | BW. Pink | 150 | 34 | 34 | 3 | 8.80% | 1 | 0 |
| pTL0047 | Jubilee | 150 | 29 | 29 | 1 | 3.40% | 1 | 0 |
| pTL0047 | Mortgage Lifter | 150 | 38 | 38 | 10 | 26.30% | 2 | 0 |
| pTL0047 | RP-BW | 150 | 80 | 80 | 33 | 41.20% | 8 | 0 |
| pTL0047 | Sunray | 150 | 33 | 33 | 8 | 24.20% | 0 | 0 |
| pTL0151 | M82 | 300 | 259 | 64 | 49 | 76.60% | 22 | 14 |
| pTL0151 | Jubilee | 250 | 276 | 132 | 30 | 22.70% | 11 | 0 |
| pTL0151 | Sunray | 250 | 136 | 100 | 30 | 30.00% | 10 | 6 |
| pTL0153 | M82 | 250 | 150 | 67 | 23 | 34.30% | 11 | 5 |
| pTL0153 | Jubilee | 250 | 200 | 69 | 23 | 33.30% | 6 | 2 |
| pTL0153 | Sunray | 250 | 125 | 101 | 35 | 34.70% | 2 | 1 |
| pTL0153 | Amana Orange | 300 | 99 | 80 | 17 | 21.30% | 30 | 6 |
| pTL0153 | BW. Pink | 250 | 97 | 52 | 26 | 50.00% | 9 | 0 |

[a]Number of explants that were suitable for infection (e.g. not damaged/contaminated).
[b] Percentage of # shoots observed divided by total infected explants.

| **Table S5. Two proportion Z-test test on transformation efficiency between pTL151 and pTL0047** | | | | | |
|---|---|---|---|---|---|
| **Genotype** | **pTL0047** | **pTL0151** | **pTL0153** | **Two proportion Z-test P-value (pTL0047 vs pTL0153)** | **Two proportion Z-test P-value (pTL0151 vs pTL0153)** |
| M82 | 29/86 | 49/64 | 23/67 | 1 (ns) | 0.000001*** |
| Jubilee | 1/29 | 30/132 | 23/69 | 0.0039*** | 0.105168 (ns) |
| Sunray | 8/33 | 30/100 | 35/101 | 0.3694 (ns) | 0.480689 (ns) |

[a]Number of shoots divided by total functional explants.

| **Table S6. Comparing timeline for transformation between pTL0151 and pTL153** | | | | | | |
|---|---|---|---|---|---|---|
| **Genotype** | **Days to rooting media (pTL151)** | **Days to rooting media (pTL0153)** | **Difference (days)** | **Days to transplanting (pTL0151)** | **Days to transplanting (pTL0153)** | **Difference (days)** |
| M82 | 62 | 64 | 2 | 19 | 11 | 8 |
| Jubilee | 78 | 53 | 25 | NA[a] | 26 | NA |
| Sunray | 61 | 56 | 5 | 15 | 10 | 5 |

[a]Rooted plants were lost due to contamination in glass jars and did not recover properly for transplanting.

**Figure S1: Vector Architecture for plasmids pTL0047, pTL0151, and pTL0153.** Vector architecture showing the expression cassettes contained within the left (LB) and right (RB) borders**.** pTL0047 contains the genes *NPTII* and *Cas9*. pTL0151 contains four guide RNAs (gRNAs) targeting the coding sequences of the genes *SP5G* and *Br*. pTL0153 was built by adding the gene *GRF4-GF1* and also contains the same gRNAs as pTL0151.

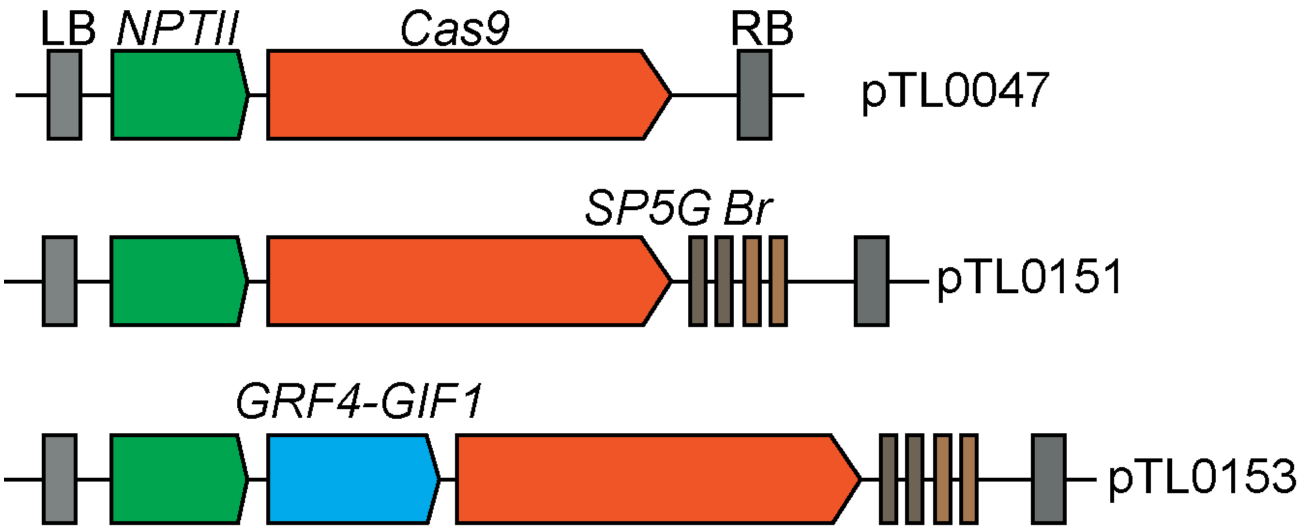

**Figure S2: Original gel electrophoresis showing the results from the genotyping by PCR performed on target genes *SP5G* and *BR*.** The first gel picture represents the amplification of the *SP5G* target sites in selected T0 plants. The second gel represents the same samples, this time showing the amplification of the *Br* target sites.

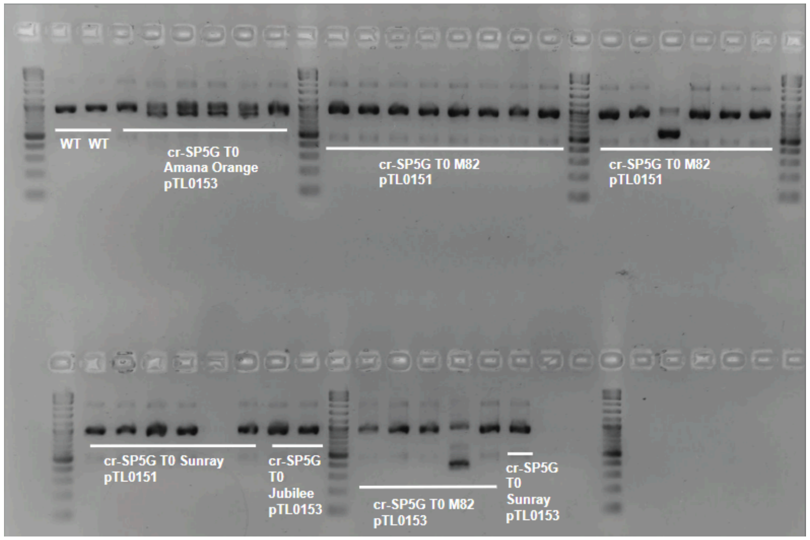


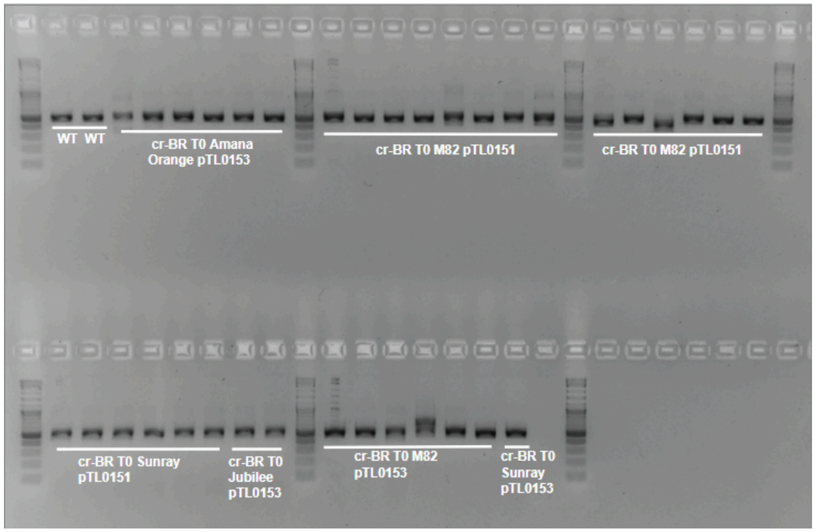
